# Software for the analysis and visualization of deep mutational scanning data

**DOI:** 10.1101/013623

**Authors:** Jesse D. Bloom

**Author notes:** Full list of author information is available at the end of the article.

## Abstract

**Background:** Deep mutational scanning is a technique to estimate the impacts of mutations on a gene by using deep sequencing to count mutations in a library of variants before and after imposing a functional selection. The impacts of mutations must be inferred from changes in their counts after selection.

**Results:** I describe a software package, `dms_tools`, to infer the impacts of mutations from deep mutational scanning data using a likelihood-based treatment of the mutation counts. I show that `dms_tools` yields more accurate inferences on simulated data than simply calculating ratios of counts pre-and post-selection. Using `dms_tools`, one can infer the preference of each site for each amino acid given a single selection pressure, or assess the extent to which these preferences change under different selection pressures. The preferences and their changes can be intuitively visualized with sequence-logo-style plots created using an extension to `weblogo`.

**Conclusions:** `dms_tools` implements a statistically principled approach for the analysis and subsequent visualization of deep mutational scanning data.

## Background

Deep mutational scanning is a high-throughput experimental technique to assess the impacts of mutations on a protein-coding gene [1]. Figure 1 shows a schematic of deep mutational scanning. A gene is mutagenized, and the library of resulting variants is introduced into cells or viruses, which are then subjected to an experimental selection that enriches for functional variants and depletes non-functional ones. Deep sequencing of the variants pre-and post-selection provides information about the functional impacts of mutations. Since the original description of deep mutational scanning by Fowler *et al* [2], the technique has been applied to a wide range of genes [3, 4, 5, 6, 7, 8, 9, 10, 11, 12, 13, 14, 17], both to measure mutational tolerance given a single selection pressure as in Figure 1A, or to identify mutations that have different effects under alternative selections as in Figure 1B. New techniques to create comprehensive codon-mutant libraries of genes make it possible to profile all amino-acid mutations [15, 16, 8, 9, 10, 17], while new techniques for targeted mutagenesis of mammalian genomes enable deep mutational scanning to be applied across the biological spectrum from viruses and bacteria to human cells [18].

**Figure 1.**
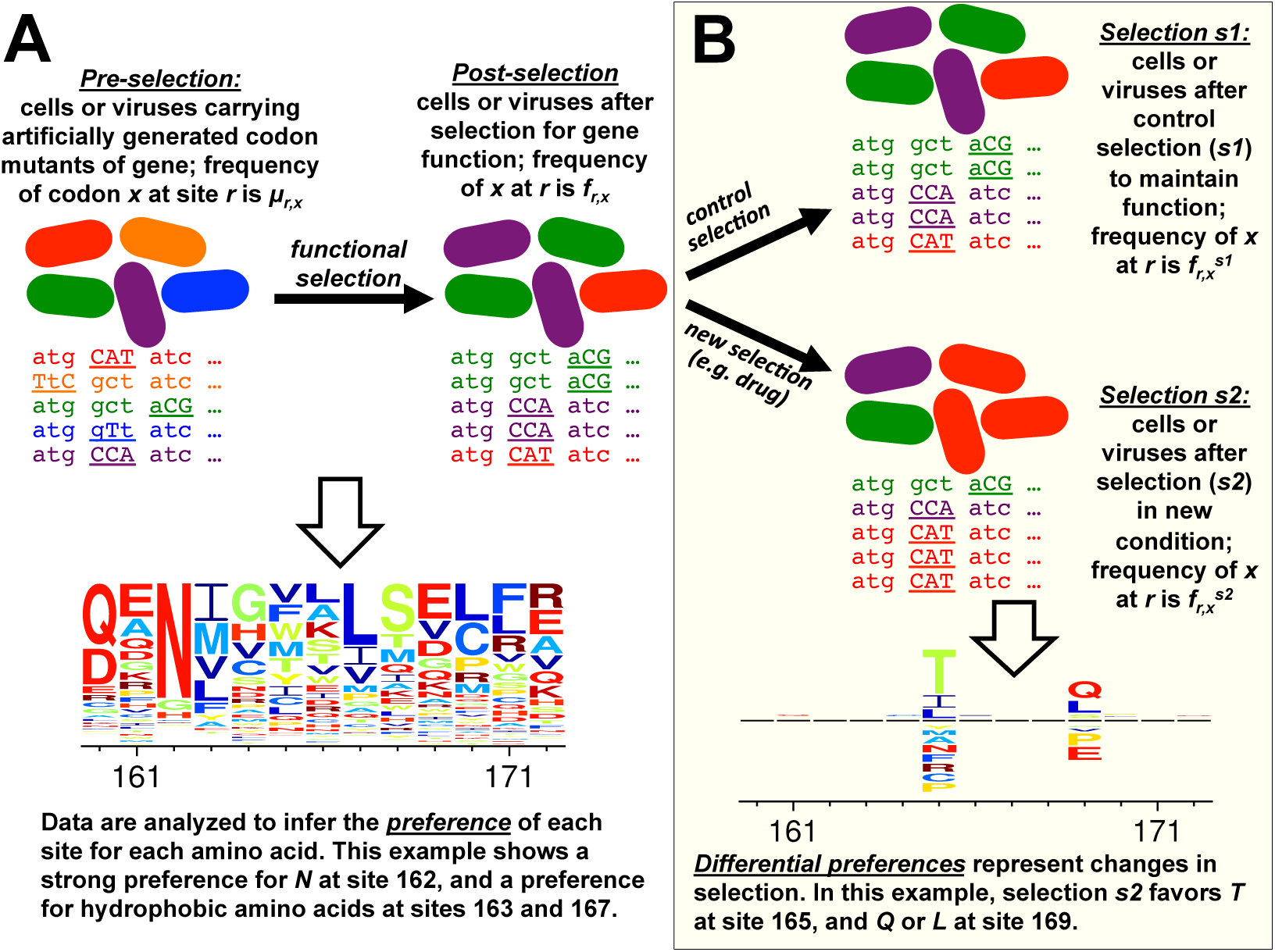
A deep mutational scanning experiment. **(A)** A gene is mutagenized to create a library that contains all single codon mutations. The mutant library is introduced into cells or viruses and subjected to a functional selection that enriches beneficial mutations and depletes deleterious ones. Deep sequencing is used to count mutations in a sample of the variants present pre-and post-selection. Using `dms_tools`, the data can be analyzed to infer the “preference” of each site for each amino acid; in the visualization, letter heights are proportional to the preference for that amino acid. **(B)** The experiment can be extended by subjecting the library of functional variants to two different selection pressures, and using deep sequencing to assess which variants are favored in one condition versus the other. Using `dms_tools`, the data can be analyzed to infer the “differential preference” of each site for each amino acid in the alternative selection *s*2 versus the control selection *s*1; in the visualization, letter heights above or below the line are proportional to the differential preference for or against that amino acid.

A key component of deep mutational scanning is analysis of the data: First, raw reads from the deep sequencing must be processed to count mutations pre-and post-selection. Next, the biological effects of mutations must be inferred from these counts. The first task of processing the reads is idiosyncratic to the specific sequencing strategy used. But the second task of inferring mutational effects from sequencing counts is amenable to more general algorithms. However, only a few such algorithms have been described [19, 20]. Here I present user-friendly software, `dms_tools`, that infers mutational effects from sequencing counts. Before describing the algorithms implemented in `dms_tools` and illustrating its use on existing and simulated data, I first discuss issues associated with inferring mutational effects from sequencing counts.

### The nature of deep mutational scanning data

The data consist of counts of variants pre-and post-selection. The approach presented here treats each site in the gene separately, ignoring epistatic coupling among mutations. This aspect of the approach should not be construed as a suggestion that interactions among mutations are unimportant; indeed, several studies have used deep mutational scanning to examine pairwise epistasis [14, 21, 22], and techniques have been described to obtain linkage between distant sites [23, 24]. However, the exploding combinatorics of multiple mutations (a 500-residue protein has only 19 × 500 ≈ 10^4^ single mutants, but 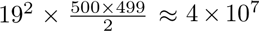 double mutants and 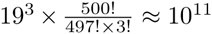 triple mutants) make it currently plausible to comprehensively characterize only single mutations to all but the shortest genes. Treating sites independently is therefore not a major limitation for most current datasets. Eventually the approach here might be extended to include coupling among mutations.

The data for each site *r* is characterized by the sequencing *depth* (total number of counts); let 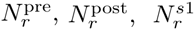, and 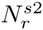 denote the depth at *r* for each of the four libraries in Figure 1 (pre-selection, post-selection, selection *s*1, and selection *s*2). Typical depths for current experiments are *N* ~ 10^6^. Denote the counts of character *x* (characters might be nucleotides, amino acids, or codons) at *r* as 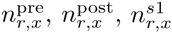, and 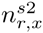. The values of *n*_*r*,*x*_ for characters *x* that differ from the wildtype identity wt (*r*) depend on both the depth *N* and the average per-site mutation rate 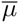. Since the mutations are intentionally introduced into the mutant library by the experimentalist, in principle 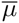 could have any value. But typically, deep mutational scanning experiments aim to introduce about one mutation per gene to avoid filling the mutant library with highly mutated genes – so the average mutation rate is usually 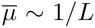 where *L* is the length of the gene. Therefore, if a 500-codon gene is sequenced at depth *N* ~ 10^6^, we expect 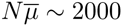 counts of non-wildtype codons at each site. Since there are 63 mutant codons, the average pre-selection counts for a mutation to a specific *x* ≠ wt (*r*) will be 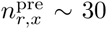, with counts for most mutations deviating from this average due to biases in creation of the mutant library and randomness in which molecules are sequenced. Counts in the post-selection libraries will further deviate from this average due to selection. Therefore, even at depths *N* ~ 10^6^, the actual counts of most mutations will be quite modest.

The rest of this paper assumes that the sequencing depth is less than the number of unique molecules in the mutant library, such that the deep sequencing randomly subsamples the set of molecules. If this assumption is false (i.e. if the number of unique molecules is substantially less than the sequencing depth), then the accuracy of inferences about mutational effects will be fundamentally limited by this aspect of the experimental design. Properly done experiments should quantify the number of unique molecules in the library so that it is obvious whether this assumption holds. In the absence of such information, the analysis can be repeated using only a random fraction of the deep sequencing data to assess whether inferences are limited by sequencing depth or the underlying molecular diversity in the mutant library.

### The goal: inferring site-specific amino-acid preferences

The goal is to estimate the effects of mutations from changes in their counts after selection. Let *μ*_*r*,*x*_, *f*_*r*,*x*_, 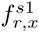, and 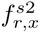 denote the *true* frequencies at site *r* of all mutant characters *x* ≠ wt (*r*) that would be observed for the four libraries in Figure 1 if we sampled at infinite depth in both the actual experiment and the sequencing. The definition of these frequencies for the wildtype character wt (*r*) depends on how the mutant library is constructed. If the mutant library is constructed so that there is a Poisson distribution of the number of mutations per gene (as is the case for error-prone PCR or the codon-mutagenesis in [9, 11]), then *μ_r_*_,wt(*r*)_, *f_r_*_,wt(*r*)_, 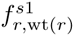, and 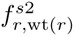 are defined as for all other characters *x*, and denote the frequencies of wt(*r*) at site *r* that would be observed if sampling at infinite depth. The reason we can make this definition for libraries containing genes with Poisson-distributed numbers of mutations is that for any reasonable-length gene (*L* ≫ 1), the marginal distribution of the number of mutations in a gene is virtually unchanged by the knowledge that there is a mutation at site *r*. On the other hand, if the mutant library is constructed so that there is exactly zero or one mutation per gene (as in [8, 10, 17]), then the marginal distribution of the total number of mutations in a gene is changed by the knowledge that there is a mutation at *r*. In this case, the wildtype-character frequencies *μ_r_*_,wt(*r*)_, *f_r_*_,wt(*r*)_, 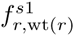, and 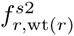 are correctly defined as the frequency of unmutated genes in the library, and the counts 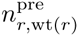, etc. are defined as the number of reads at *r* attributable to unmutated genes. In this case, measurement of these counts requires sequencing with linkage as in [23, 24, 17]. The proper analysis of libraries containing only unmutated and singly mutated clones sequenced without linkage is beyond the scope of this paper.

If we knew the frequencies *μ*_*r*,*x*_, *f*_*r*,*x*_, 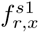, and 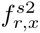, we could calculate parameters that reflect the effects of mutations. One parameter that characterizes the effect of mutating *r* from wt (*r*) to *x* for the experiment in Figure 1A is the *enrichment ratio*, which is the relative frequency of mutations to *x* after selection versus before selection:

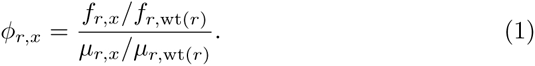

Beneficial mutations have *ϕ*_*r*,*x*_ > 1, while deleterious ones have *ϕ*_*r*,*x*_ < 1. A related parameter is the *preference π*_*r*,*x*_ of *r* for *x*. At each site, the preferences are simply the enrichment ratios rescaled to sum to one:

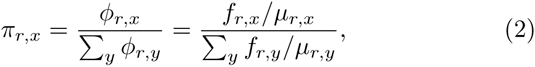

or equivalently

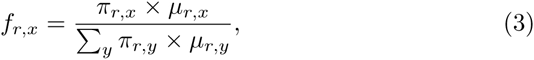

where *y* is summed over all character identities (all nucleotides, codons, or amino acids). The preferences can be intuitively visualized (Figure 1A) and interpreted as the equilibrium frequencies in substitution models for gene evolution [9, 25] (after accounting for uneven mutational rates [26, 27]).

### The challenge of statistical inference from finite counts

Equations 1 and 2 are in terms of the true frequencies *μ*_*r*,*x*_, *f*_*r*,*x*_, etc. But in practice, we only observe the counts in the finite sample of sequenced molecules. The computational challenge is to estimate the preferences (or enrichment ratios) from these counts.

The most naive approach is to simply substitute the counts for the frequencies, replacing Equation 1 with

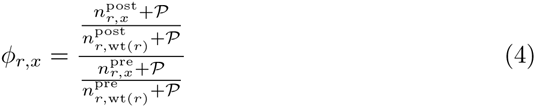

where 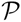 (often chosen to be one) is a pseudocount added to each count to avoid ratios of zero or infinity.

However, Equation 4 involves ratios of counts with values ~ 10 to 100 – and as originally noted by Karl Pearson [28, 29], ratios estimated from finite counts are statistically biased, with the bias increasing as the magnitude of the counts decrease. This bias can propagate into subsequent analyses, since many statistical tests assume symmetric errors. The problems caused by biases in uncorrected ratios have been noted even in applications such as isotope-ratio mass spectrometry [30] and fluorescent imaging [31], where the counts usually far exceed those in deep mutational scanning.

Taking ratios also abrogates our ability to use the magnitude of the counts to assess our certainty about conclusions. For instance, imagine that at a fixed depth, the counts of a mutation increase from a pre-selection value of 5 to a post-selection value of 10. While this doubling suggests that the mutation might be beneficial, the small counts make us somewhat uncertain of this conclusion. But if the counts increased from 20 to 40 we would be substantially more certain, and if they increased from 100 to 200 we would be quite sure. So only by an explicit statistical treatment of the counts can we fully leverage the data.

Here I describe a software package, `dms_tools`, that infers mutational effects in a Bayesian framework using a likelihood-based treatment of the counts. This software can be used to infer and visualize site-specific preferences from experiments like Figure 1A, and to infer and visualize differences in preferences under alternative selections from experiments like Figure 1B.

## Implementation and Results

### Algorithm to infer site-specific preferences

`dms_tools` uses a Bayesian approach to infer site-specific preferences from experiments like those in Figure 1A. The algorithm calculates the likelihoods of the counts given the unknown preferences and mutation / error rates, placing plausible priors over these unknown parameters. The priors correspond to the assumption that all possible identities (e.g. amino acids) have equal preferences, and that the mutation and error rates for each site are equal to the overall average for the gene. MCMC is used to calculate the posterior probability of the preferences given the counts.

This algorithm is a slight modification of that in the *Methods* of [9]; here the algorithm is described anew to explain the implementation in `dms_tools`.

#### Optional controls to quantify error rates

Some sequencing reads that report a mutation may actually reflect an error introduced during sequencing or PCR rather than an actual mutation that experienced selection. Errors can be quantified by sequencing an unmutated gene, so that any counts at *r* of *x* ≠ wt(*r*) for this control reflect errors. In some cases (e.g. sequencing an RNA virus where the post-selection libraries must be reverse-transcribed), error rates for the pre-and post-selection libraries may differ and so be described by different controls. Let 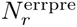 and 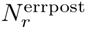 be the depth and 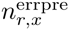 and 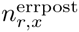 be the counts of *x* in the pre-selection and post-selection error controls, respectively. Define *ϵ*_*r*,*x*_ and *ρ*_*r*,*x*_ to be the true frequencies of errors at *r* from wt (*r*) to *x* in the pre-and post-selection controls, respectively.

#### Likelihoods of observing specific mutational counts

Define vectors of the counts and frequencies for all characters at each site *r*, i.e. 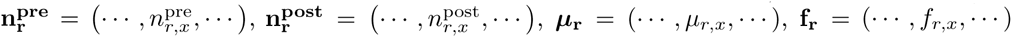, etc. Also define 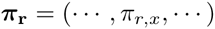 of the preferences for each *r*, noting that Equation 3 implies 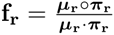 where ○ is the Hadamard product.

The likelihoods of some specific set of counts are:

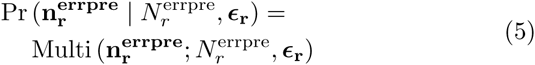

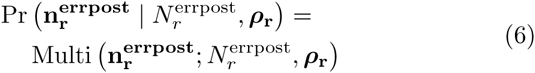

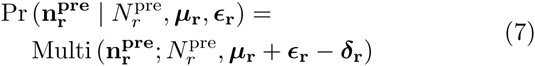

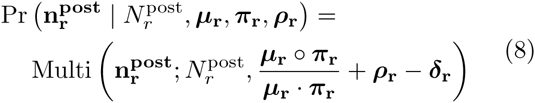

where Multi is the multinomial distribution, 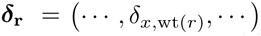 is a vector with the element corresponding to wt (*r*) equal to one and all other elements zero (*δ_xy_* is the Kronecker delta), and we have assumed that the probability that a site experiences both a mutation and an error is negligibly small.

#### Priors over the unknown parameters

We specify Dirichlet priors over the parameter vectors:

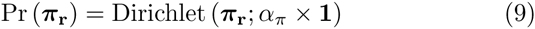

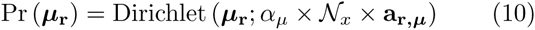

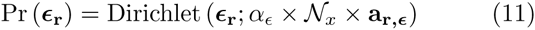

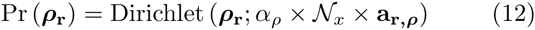

where **1** is a vector of ones, 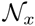 is the number of characters (64 for codons, 20 for amino acids, 4 for nucleotides), the *α*’s are scalar concentration parameters > 0 (by default `dms_tools` sets the *α*’s to one). For codons, the error rate depends on the number of nucleotides being changed. The average error rates 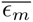 and 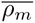 for codon mutations with *m* nucleotide changes are estimated as

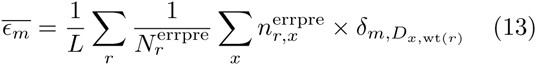

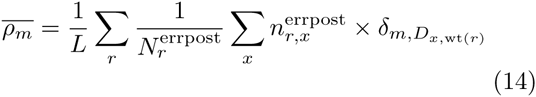

where *D_x_*_,wt(*r*)_ is the number of nucleotide differences between *x* and wt (*r*). Given these definitions,

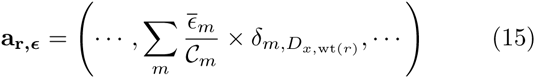

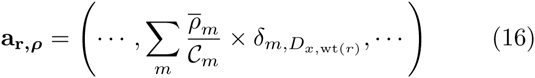

where 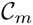 is the number of mutant characters with *m* changes relative to wildtype (for nucleotides 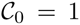 and 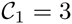; for codons 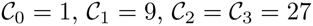).

Our prior assumption is that the mutagenesis introduces all mutant characters at equal frequency (this assumption is only plausible for codons if the mutagenesis is at the codon level as in [15, 16, 8, 9, 10, 17]; if mutations are made at the nucleotide level such as by error-prone PCR then characters should be defined as nucleotides). The average per-site mutagenesis rate is estimated as

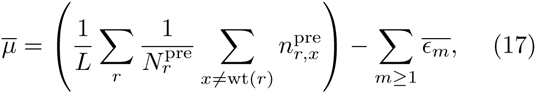

so that

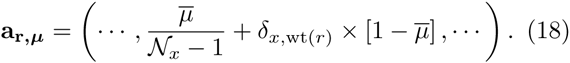

#### Character types: nucleotides, amino acids, or codons

`dms_tools` allows four possibilities for the type of character for the counts and preferences. The first three possibilities are simple: the counts and preferences can both be for any of nucleotides, amino acids, or codons.

The fourth possibility is that the counts are for codons, but the preferences for amino acids. In this case, define a function mapping codons to amino acids,

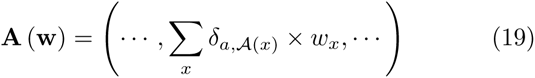

where **w** is a 64-element vector of codons *x*, **A** (**w**) is a 20-or 21-element (depending on the treatment of stop codons) vector of amino acids *a*, and 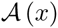 is the amino acid encoded by *x*. The prior over the preferences ***π*_r_** is still a symmetric Dirichlet (now only of length 20 or 21), but the priors for ***μ*_r_**, ***ϵ*_r_**, and ***ρ*_r_** are now Dirichlets parameterized by **A** (**a_r,*μ*_**), **A** (**a_r,_*_ϵ_***) and **A** (**a_r,*ρ*_**) rather than **a_r,*μ*_**, **a_r,_*_ϵ_***, and **a_r,*ρ*_**. The likelihoods are computed in terms of these transformed vectors after similarly transforming the counts to 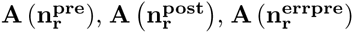 and 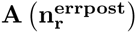.

#### Implementation

The program `dms_inferprefs` in the `dms_tools` package infers the preferences by using `pystan` [32] to perform MCMC over the posterior defined by the product of the likelihoods and priors in Equations 5, 6, 7, 8, 9, 10, 11, and 12. The program runs four chains from different initial values, and checks for convergence by ensuring that the Gelman-Rubin statistic 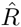 [33] is < 1.1 and the effective sample size is > 100; the number of MCMC iterations is increased until convergence is achieved. The program `dms_logoplot` in the `dms_tools` package visualizes the posterior mean preferences via an extension to `weblogo` [34]. The program `dms_merge` can be used to average preferences inferred from different experimental replicates that have individually been analyzed by `dms_inferprefs`, and the program `dms_correlate` can be used to compute the correlations among inferences from different replicates.

### Inferring preferences with dms_tools

#### Application to actual datasets

Figures 2 and 3 illustrate application of `dms_tools` to two existing datasets [10, 11]. The programs require as input only simple text files listing the counts of each character identity at each site. As the figures show, the `dms_inferprefs` and `dms_logoplot` programs can process these input files to infer and visualize the preferences with a few simple commands. Error controls can be included when available (they are not for Figure 2, but are for Figure 3). The runtime for the MCMC depends on the gene length and character type (codons are slowest, nucleotides fastest) – but if the inference is parallelized across multiple CPUs (using the `--ncpus` option of `dms_inferprefs`), the inference should take no more than a few hours. As shown in Figures 2 and 3, the visualizations can overlay information about protein structure onto the preferences.

**Figure 2.**
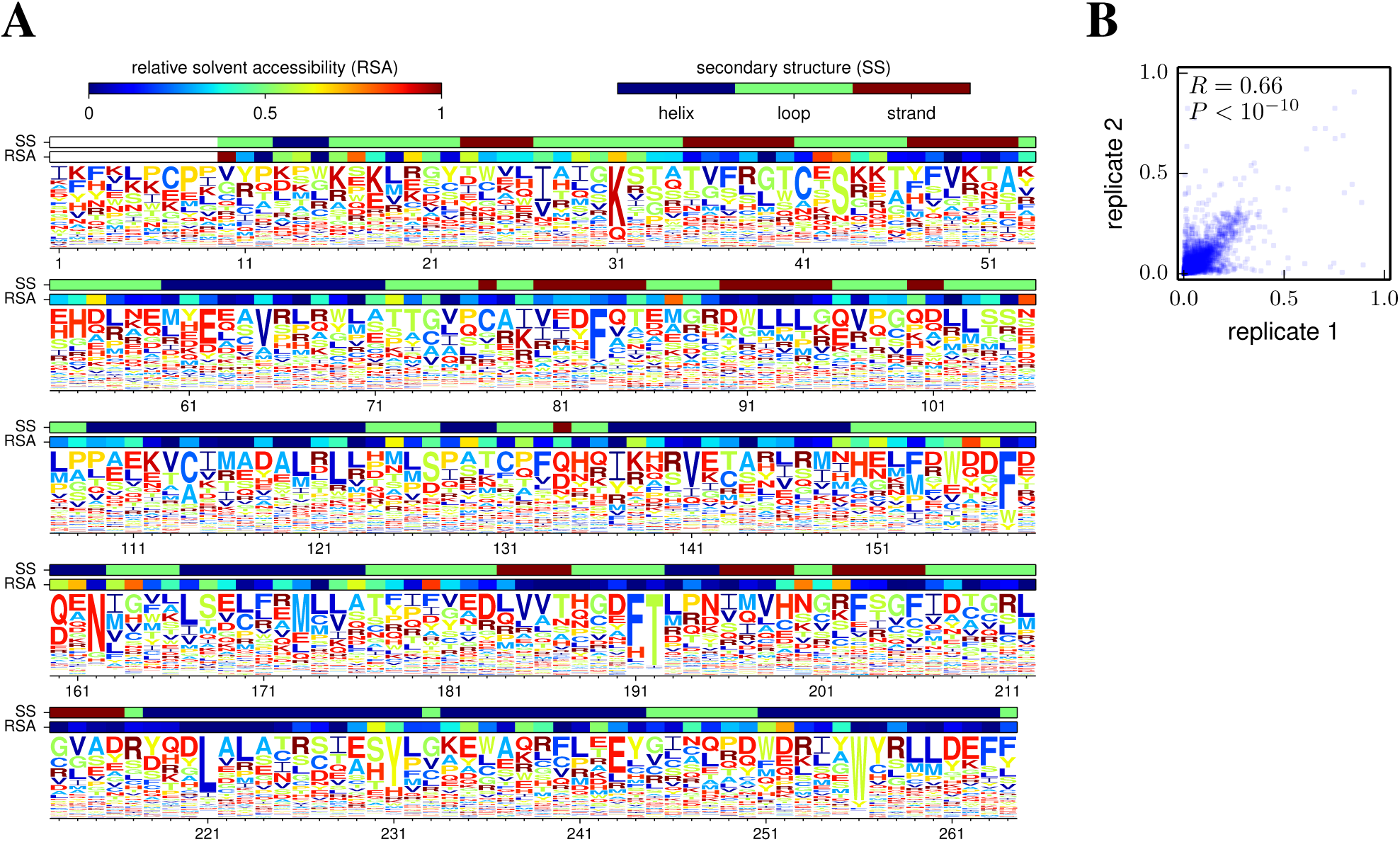
Site-specific preferences from deep mutational scanning of a Tn5 transposon. Melnikov *et al* [10] performed deep mutational scanning on a Tn5 transposon using kanamycin selection, and reported the counts of amino-acid mutations for two biological replicates of the experiment. Here I have used `dms_tools` to infer the preferences. **(A)** Visualization of the preferences averaged across the two replicates. **(B)** Correlation between the preferences inferred from each of the two replicates. Given files containing the mutation counts, the plots can be generated as `logoplot.pdf` and `corr.pdf` with the following commands:

~~~
dms_inferprefs pre_counts_1.txt post_counts_1.txt prefs_1.txt --excludestop --ncpus -1 --chartype aa
dms_inferprefs pre_counts_2.txt post_counts_2.txt prefs_2.txt --excludestop --ncpus -1 --chartype aa
dms_correlate prefs_1.txt prefs_2.txt corr --name1 "replicate 1" --name2 "replicate 2" --corr_on_plot
dms_merge prefs.txt average prefs_1.txt prefs_2.txt
dms_logoplot prefs.txt logoplot.pdf --nperline 53 --overlay1 RSAs.txt RSA "relative solvent accessibility”
     --overlay2 SSs.txt SS "secondary structure”
~~~

**Figure 3.**
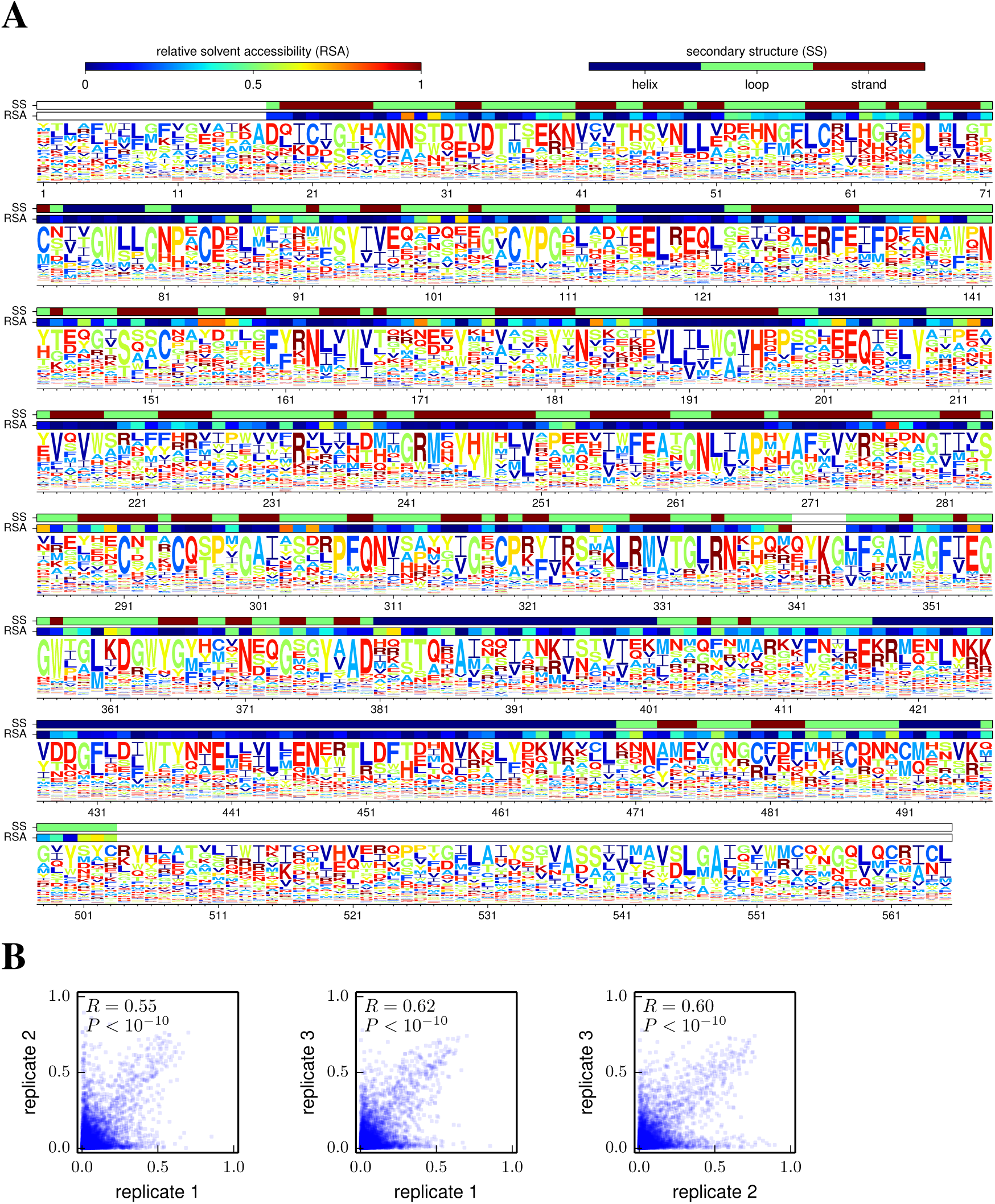
Site-specific preferences from deep mutational scanning of influenza hemagglutinin. Thyagarajan and Bloom [11] performed deep mutational scanning on influenza hemagglutinin, and reported the counts of codon mutations for three biological replicates of the experiment. Here I have used `dms_tools` to infer the preferences. **(A)** Visualization of the preferences averaged across the three replicates. **(B)** Correlations between the preferences from each pair of replicates. Given files containing the mutation counts, the plots can be generated as `logoplot.pdf`, `corr 1 2.pdf`, `corr 1 3.pdf`, and `corr 2 3.pdf` with the following commands:

~~~
dms_inferprefs mutDNA_1.txt mutvirus_1.txt prefs_1.txt --errpre DNA_1.txt --errpost virus_1.txt --ncpus -1
dms_inferprefs mutDNA_2.txt mutvirus_2.txt prefs_2.txt --errpre DNA_2.txt --errpost virus_2.txt --ncpus -1
dms_inferprefs mutDNA_3.txt mutvirus_3.txt prefs_3.txt --errpre DNA_3.txt --errpost virus_3.txt --ncpus -1
dms_correlate prefs_1.txt prefs_2.txt corr_1_2 --name1 "replicate 1" --name2 "replicate 2" --corr_on_plot
dms_correlate prefs_1.txt prefs_3.txt corr_1_3 --name1 "replicate 1" --name2 "replicate 3" --corr_on_plot
dms_correlate prefs_2.txt prefs_3.txt corr_2_3 --name1 "replicate 2" --name2 "replicate 3" --corr_on_plot
dms_merge prefs.txt average prefs_1.txt prefs_2.txt prefs_3.txt
dms_logoplot prefs.txt logoplot.pdf --nperline 71 --overlay1 RSAs.txt RSA "relative solvent accessibility”
     --overlay2 SSs.txt SS "secondary structure" --excludestop
~~~ Note that unlike in Figure 2, no `--chartype` option is specified since the `dms_inferprefs` default is already `codon_to_aa`.

Figures 2 and 3 also illustrate use of `dms_correlate` to assess the correlation between preferences inferred from different biological replicates [35] of the experiment. The inclusion and analysis of such replicates is the only sure way to fully assess the sources of noise associated with deep mutational scanning.

#### Testing on simulated data

To test the accuracy of preference-inference by `dms_tools`, I simulated deep mutational scanning counts using the preferences in Figure 2, both with and without errors quantified by appropriate controls. Importantly, the error and mutation rates for these simulations were *not* uniform across sites and characters, but were simulated to have a level of unevenness comparable to that observed in real experiments. I then used `dms_tools` to infer preferences from the simulated data, and also made similar inferences using simple ratio estimation (Equation 4). Figure 4 shows the inferred preferences versus the actual values used to simulate the data. For simulations with mutation counts (quantified by the product 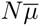 of the depth and average per-site mutation rate) ~ 1000 to 2000, the inferences are quite accurate. Inferences made by `dms_tools` are always more accurate than those obtained by simply taking ratios of mutation counts.

**Figure 4.**
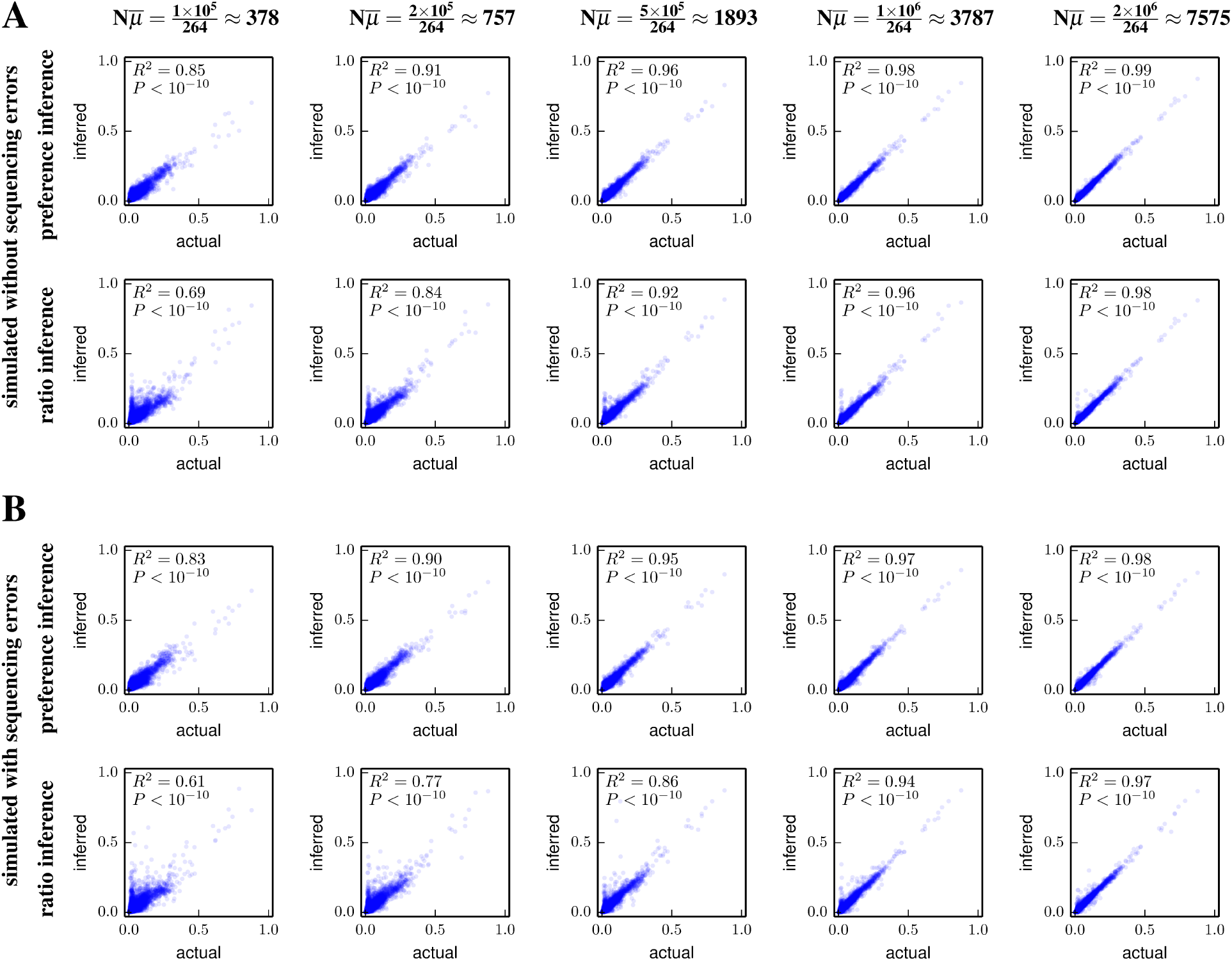
Accuracy of preference inference on simulated data. Deep mutational scanning counts were simulated using the preferences in Figure 2A and realistic mutation and error rates that were uneven across sites and characters as in actual experiments. The simulations were done **(A)** without or **(B)** with sequencing errors quantified by control libraries. Plots show the correlation between the actual and inferred preferences as a function of the product of the sequencing depth *N* and the average per-site mutation rate 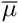; real experiments typically have 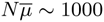 to 2000 depending on the sequencing depth and gene length. Preferences are inferred using the full algorithm in `dms_tools` (top panels) or by simply calculating ratios of counts (bottom panels) using Equation 4 and its logical extension to include errors, both with a pseudocount of one. The `dms_tools` inferences are more accurate than the simple ratio estimation, with both methods converging to the actual values with increasing 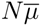. Given files with the mutation counts, the plots in this figure can be generated as `prefs corr.pdf` and `ratio corr.pdf` with commands such as:

~~~
dms_inferprefs pre.txt post.txt inferred_prefs.txt --ncpus -1
dms_inferprefs pre.text post.text ratio_prefs.txt --ratio_estimation 1
dms_correlate actual_prefs.txt inferred_prefs.txt prefs_corr --name1 "actual" --name2 "inferred”
     --corr_on_plot --r2
dms_correlate actual_prefs.txt ratio_prefs.txt ratio_corr --name1 "actual" --name2 "inferred" --corr_on_plot
     --r2
~~~

#### Is the Bayesian inference worthwhile?

The foregoing sections explain why the Bayesian inference of preferences implemented in `dms_tools` is conceptually preferable to estimating mutational effects via direct ratio estimation using Equation 4. However, do the practical benefits of this Bayesian inference justify its increased complexity? The simulations in the previous section show that the Bayesian inference is more accurate, but in the absence of background errors (Figure 4A) the magnitude of the improvement becomes negligible once the mutation counts per site 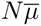 start to exceed ~ 10^3^. When there is a need to correct for background errors (Figure 4B), meaningful benefits of the Bayesian inference over enrichment ratios extend to somewhat higher sequencing depths. Overall, it appears that Bayesian inference will always perform as well or better than ratio estimation, but that the tangible benefit becomes negligible at high sequencing depth. In that case, the user will have to decide if the increased computational runtime and complexity of the Bayesian inference is worth a small improvement in accuracy. Simpler ratio estimation can be performed using the `--ratio_estimation` option of `dms_inferprefs` or using an alternative program such as `Enrich` [19]. When applying ratio estimation to data where some mutations have low counts, it is important to include pseudocounts (denoted by 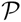 in Equation 4) as a form of regularization to avoid estimating excessively high or low preferences at sites with limited counts.

### Algorithm to infer differential preferences

As shown in Figure 1B, a useful extension to the experiment in Figure 1A is to subject the functional variants to two different selection pressures to identify mutations favored by one pressure versus the other. While this experiment could in principle by analyzed by simply comparing the initial unselected mutants to the final variants after the two alternative selections, this approach is non-ideal. In experiments like Figure 1A, many mutations are enriched or depleted to some extent by selection, since a large fraction of random mutations affect protein function [36, 37, 38, 39, 40]. Therefore, the assumption that all mutations are equally tolerated (i.e. the preferences for a site are all equal, or the enrichment ratios are all one) is not a plausible null hypothesis for Figure 1A. For this reason, `dms_tools` simply infers the preferences given a uniform Dirichlet prior rather than trying to pinpoint some subset of sites with unequal preferences.

But in Figure 1B, the assumption that most mutations will be similarly selected is a plausible null hypothesis, since we expect alternative selections to have markedly different effects on only a small subset of mutations (typically, major constraints related to protein folding and stability will be conserved across different selections on the same protein). Therefore, `dms_tools` uses a different algorithm to infer the differential preferences under the two selections. This algorithm combines a prior that mildly favors differential preferences of zero with a likelihood-based analysis of the mutation counts to estimate posterior probabilities over the differential preferences.

#### Definition of the differential preferences

Given an experiment like Figure 1B, let 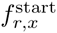 be the true frequency of character *x* at site *r* in the starting library (equivalent to the frequency 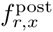 in the figure), and let 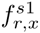 and 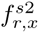 be the frequencies after selections *s*1 and *s*2, respectively. The differential preference Δ*π*_*r*,*x*_ for *x* at *r* in *s*2 versus *s*1 is defined by:

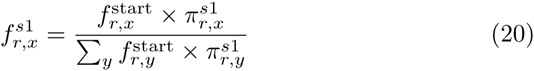

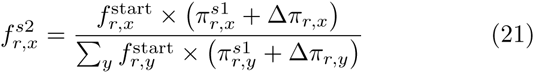

where 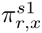 is the “control preference" and is treated as a nuisance parameter, and we have the constraints

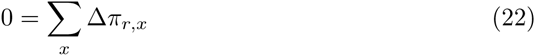

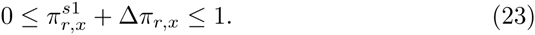

If there is no difference in the effect of *x* at *r* between selections *s*1 and *s*2, then Δ*π*_*r*,*x*_ = 0. If *x* at *r* is more preferred by *s*2 than *s*1, then Δ*π*_*r*,*x*_ > 0; conversely if *x* at *r* is more preferred by *s*1 than *s*2, then Δ*π*_*r*,*x*_ < 0 (see Figure 5A).

**Figure 5.**
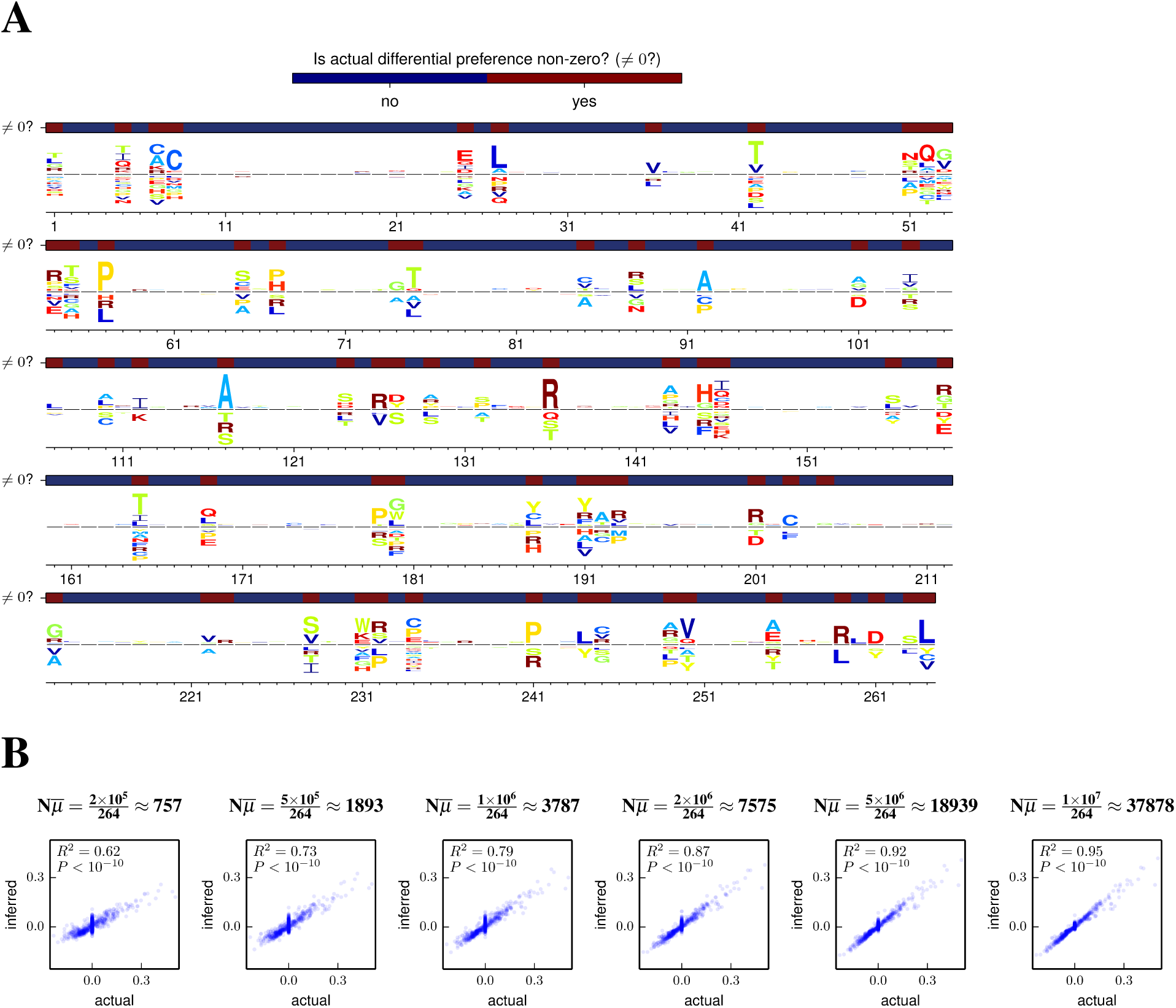
Inference of differential preferences on simulated data. To illustrate and test the inference of differential preferences, the experiment in Figure 1B was simulated at the codon level starting with the post-selection library that yielded the preferences in Figure 2. In the simulations, 20% of sites had different preferences between the control and alternative selection. **(A)**, `dms_tools` was used to infer the differential preferences from the data simulated at *N* = 10^7^, and the resulting inferences were visualized. The overlay bars indicate which sites had non-zero differential preferences in the simulation. **(B)** The correlations between the inferred and actual differential preferences as a function of 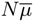 show that the inferred values converge to the true ones. Given files with the mutation counts, the plots in this figure can be generated as `logoplot.pdf` and `corr.pdf` with the following commands:

~~~
dms_inferdiffprefs start.txt s1.txt s2.txt diffprefs.txt --ncpus -1
dms_logoplot diffprefs.txt logoplot.pdf --nperline 53 --overlay1 actually_nonzero.txt "$\ne 0$?" "Is actual
     differential preference non-zero?" --diffprefheight 0.45
dms_correlate actual_diffprefs.txt diffprefs.txt corr --name1 "actual" --name2 "inferred" --corr_on_plot --r2
~~~ Note that no `--chartype` option is specified because the default for `dms_inferdiffprefs` is already `codon_to_aa`.

#### Likelihoods of observing specific mutational counts

Define vectors of the counts as 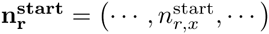 for the post-selection functional variants that are subjected to the further selections, and as 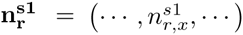 and 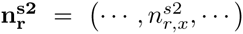 for selections *s*1 and *s*2. We again allow an error control, but now assume that the same control applies to all three libraries (since they are all sequenced after a selection), and define the counts for this control as 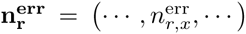; the true error frequencies are denoted by *ξ*_*r*,*x*_. Define vectors of the frequencies, errors, control preferences, and differential preferences: 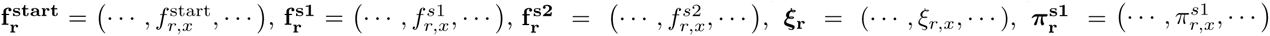, and 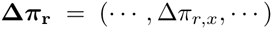. Equations 20 and 21 imply 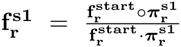 and 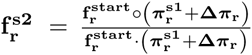.

The likelihoods of the counts will be multinomially distributed around the “true” frequencies, so

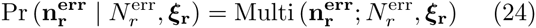

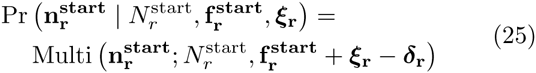

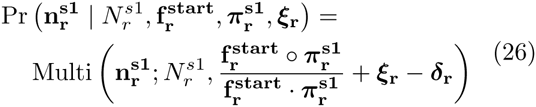

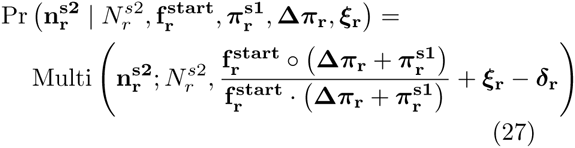

where we have assumed that the probability that a site experiences a mutation and an error in the same molecule is negligibly small.

#### Priors over the unknown parameters

We specify Dirichlet priors over the parameter vectors:

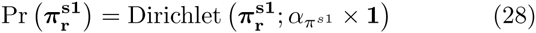

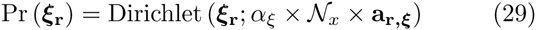

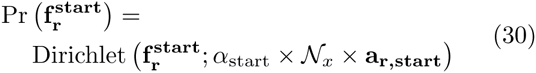

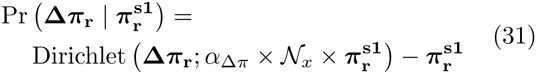

where `dms_tools` by default sets all the scalar concentration parameters (*α*’s) to one except *α*_Δ*π*_, which is set to two corresponding to a weak expectation that the Δ*π* values are close to zero. The average error rate 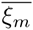 for mutations with *m* nucleotide changes is

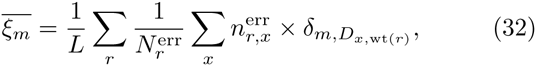

and so

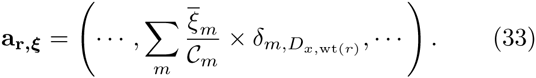

Our prior assumption is that all mutations are at equal frequency in the starting library (this assumption is unlikely to be true if the starting library has already been subjected to some selection, but we lack a rationale for a more informative prior). The average mutation frequency in the starting library is

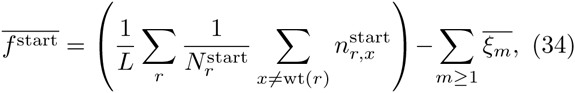

and so

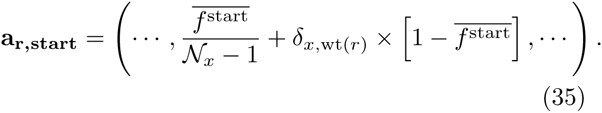

#### Implementation

The program `dms_inferdiffprefs` in the `dms_tools` package infers the differential preferences by performing MCMC over the posterior defined by the product of the likelihoods and priors in Equations 24, 25, 26, 27, 28, 29, 30, and 31. The MCMC is performed as described for the preferences, and characters can again be any of nucleotides, amino acids, or codons. The program `dms_logoplot` visualizes the posterior mean differential preferences via an extension to `weblogo` [34]. In addition, `dms_inferdiffprefs` creates text files that give the posterior probability that Δ*π*_*r*,*x*_ > 0 or < 0. These posterior probabilities are *not* corrected to account for the fact that multiple sites are typically being examined, although by default the inferences are made using the regularizing prior in Equation 31.

### Inferring differential preference with dms_tools

To test the accuracy of differential preference inference by `dms_tools`, I simulated an experiment like that in Figure 1B with the starting counts based on Melnikov *et al* ‘s actual deep mutational scanning data of a Tn5 transposon [10]. As shown by Figure 5, `dms_inferdiffprefs` accurately infers the differential preferences at typical experimental depths. The results are easily visualized with `dms_logoplot`. To provide a second illustration of differential preferences, Figure 6 shows an analysis of the data obtained by Wu *et al* when they performed an experiment like that in Figure 1B on nucleotide mutants of the influenza NS gene in the presence or absence of interferon treatment.

**Figure 6.**
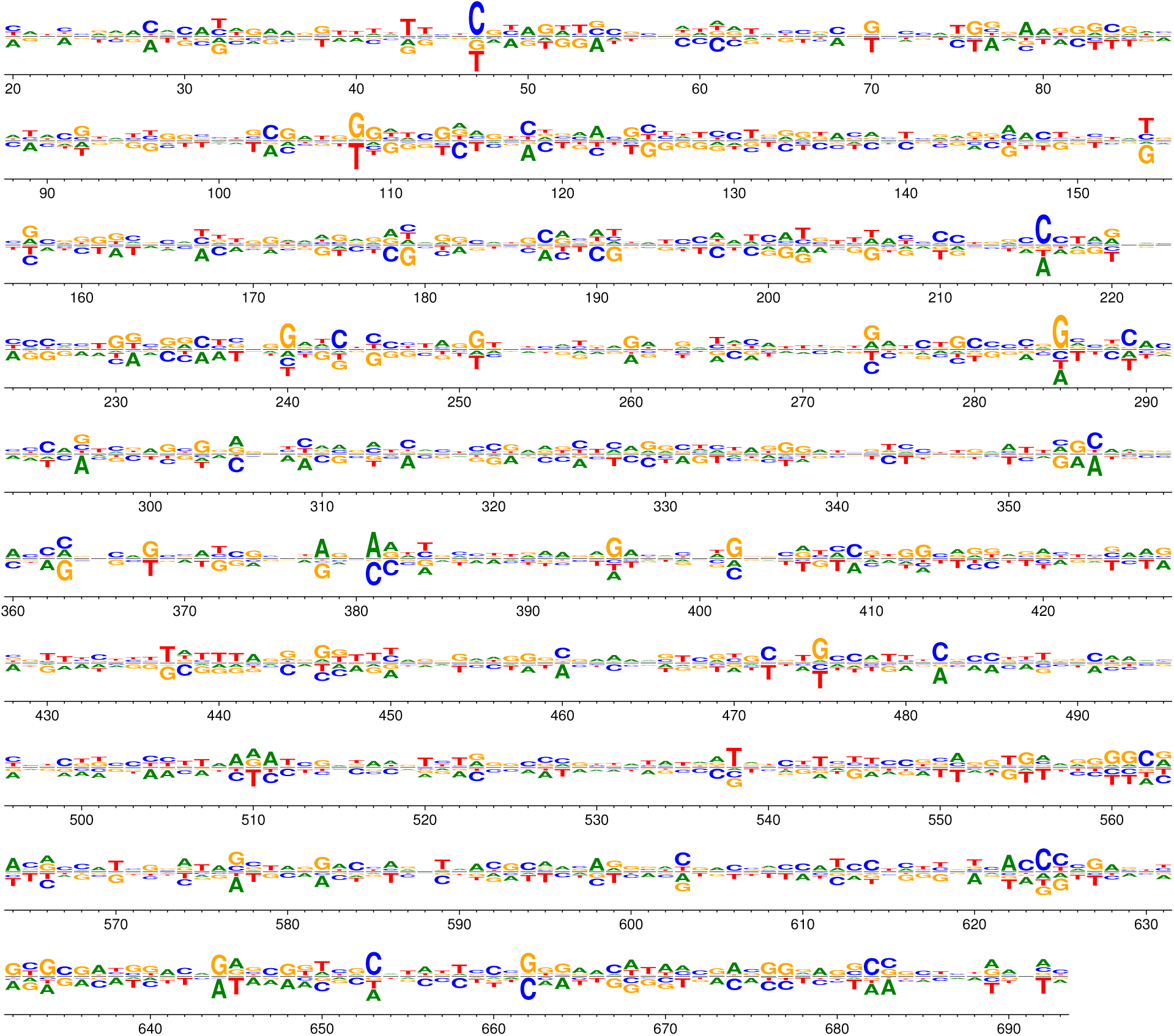
Differential preferences following selection of influenza NS1 in the presence or absence of interferon. Wu *et al* [13] generated libraries of influenza viruses carrying nucleotide mutations in the NS segment. They passaged these viruses in the presence or absence of interferon pre-treatment. Here, `dms_tools` was used to analyze and visualize the data to identify sites where different nucleotides are preferred in the presence versus the absence of interferon. Because the mutations were made at the nucleotide level, the data must also be analyzed at that level (unlike in Figures 2, 3, and 5, where codon mutagenesis means that the data can be analyzed at the amino-acid level). The plot can be generated as `logoplot.pdf` with the following commands:

~~~
dms_inferdiffprefs input.txt control.txt interferon.txt diffprefs.txt --ncpus -1 --chartype DNA
dms_logoplot diffprefs.txt logoplot.pdf --nperline 68 --diffprefheight 0.4
~~~

## Conclusions

`dms_tools` is a freely available software package that uses a statistically principled approach to analyze deep mutational scanning data. This paper shows that `dms_tools` accurately infers preferences and differential preferences from data simulated under realistic parameters. As the figures illustrate, `dms_tools` can also be applied to actual data with a few simple commands. The intuitive visualizations created by `dms_tools` assist in interpreting the results. As deep mutational scanning continues to proliferate as an experimental technique [1], `dms_tools` can be applied to analyze the data for purposes such as guiding protein engineering [3, 10], understanding sequence-structure-function relationships [4, 5, 7, 14, 21], informing the development of better evolutionary models for sequence analysis [9, 25], and probing the biology of viruses and cells [6, 8, 11, 12, 13, 18].

## Availability and requirements

- **Project name:** `dms_tools`
- **Project home page:**

- Documentation and installation instructions: http://jbloom.github.io/dms_tools/
- Source code: https://github.com/jbloom/dms_tools
- **Operating system(s):** Linux
- **Programming language:** `Python`
- **Other requirements:** `pystan`, `weblogo`
- **License:** GNU GPLv3
- **Restrictions to use by non-academics:** None

## Data and code for figures in this paper

The data and computer code used to generate the figures are in version 1.01 of the `dms_tools` source code (which is tagged on `Github` at https://github.com/jbloom/dms_tools/tree/1.0.1) in the `examples` subdirectory. The `LaTex` source for this paper is in the `paper` subdirectory.

## Author’s contributions

JDB designed the algorithms, wrote the software, performed the analyses, and wrote the paper.

## Acknowledgements

Thanks to Alec Heckert for assistance in testing `dms_tools`, to Erick Matsen for the excellent suggestion to use `pystan` for MCMC, to Nicholas Wu for providing the mutational counts data from [13], and to Orr Ashenberg and Hugh Haddox for helpful comments on the manuscript. This work was supported by the NIGMS of the NIH under grant R01GM102198.

